# Prosocial reward relates to performance in difficult speech-on-speech listening conditions

**DOI:** 10.1101/2024.10.02.616312

**Authors:** Ryan J Oakeson, Stuart Rosen, Jesca Herbert, Simon Roper, Hongyi Zhang, Sophie K Scott

## Abstract

Motivation plays an important role in a listener’s effort when in noisy environments. However, there remain gaps in the previous literature pertaining to the social psychological factors underlying motivation and listening effort. To fill these gaps, this study explored how prosociality and social reward relate to speech perception in noise (SPiN) in a group of normal hearing English speakers (n = 136; mean age: 29.6, age range: 18-68). We investigated SPiN performance and subjective listening experiences across different speech masking conditions: 1-speaker, 2-speaker, and speech-spectrum shaped noise (SSN), along with a working memory task, and questionnaires pertaining to social orientation. Results indicated a robust effect of different maskers, and that individuals who rated themselves higher in prosocial traits performed better in the masker that yielded the highest threshold out of the three conditions, the 2-speaker condition. Additionally, subjective ratings of listening effort particularly in the 1-speaker condition related to age, where older participants reported greater effort. These findings highlight prosociality and age as important social psychological factors influencing SPiN performance and listening effort, respectively, particularly in complex listening scenarios.

## II. Introduction

Perceiving speech in noisy environments is a difficult task that requires a listener to accomplish several different goals simultaneously. Not only must the listener use intention to ignore or attenuate irrelevant streams of auditory input, but they must selectively focus their auditory attention to the target speaker using cognitive effort and control (Eckert et al., 2016). This is the essence of the “cocktail party” problem (Cherry, 1953). This process leads to a complex state in which the listener is exerting substantial amounts of listening effort—often increasing mental fatigue, decreasing motivation, and having other social psychological consequences (Peelle, 2016; Pichora-Fuller et al., 2016). By understanding how the brain is capable of such selectivity may bring us steps closer to understanding the complexity of attention during cocktail party situations. However, objectively measuring and understanding the underlying economics involved in effortful listening proves to be a difficult task in-and-of itself (Eckert et al., 2016; McGarrigle et al., 2014). Following this notion, numerous researchers have been interested in the various components underlying the “cocktail party” problem and how to solve the mystery as to how the brain performs and overcomes such adversity.

Here we aim to gain insight into how social psychological factors interact with speech perception in noise (SPiN). First, definitions of “noise” will be presented in the context of language perception, examples of how motivators have been shown to modulate performance and physiology during studies of cognitive effort will be addressed, then followed by framing motivation in terms of prosociality. Lastly, psycho-social traits related to described and how they may be related to SPiN, followed by the main aims and hypotheses of the study.

### A. Speech Perception in Noise

Speech is rarely processed in silence. Fundamentally, where selective attention is key process in the cocktail party effect, masking sounds are the key issue in the cocktail party problem. Whilst the power or salience of the masking sound varies from situation-to-situation, the type of maskers in the environment plays a significant role in outcomes related to SPiN. Maskers in this case can be divided into two types, energetic/modulation and informational (Cooke, 2006; Shinn-Cunningham, 2008; Stone et al., 2012). Energetic/modulation maskers suppress or distort speech signals at the signal level whereas informational maskers encompass everything beyond the signal level, and typically consist of other speech (Kidd et al., 2008). For example, informational maskers are found to typically disrupt speech perception at higher-order perceptual and cognitive levels related to language and other implicated capacities e.g., attention and memory (Mattys et al., 2012; Rönnberg et al., 2013). Therefore, various speech masking noises may result in people exerting increased listening effort due to high demands on mental resources, resulting in the listener evaluating the costs and benefits related to mental resource allocation (Eckert et al., 2016; Pichora-Fuller et al., 2016). One way to combat these increasing demands is by motivating the listener’s effort.

### B. Motivation

Motivation has been found to play a critical role in listening performance and subjective experiences of effort and fatigue, as well as how individuals utilize their mental resources based on costs and rewards (Carolan et al., 2022; Pichora-Fuller et al., 2016; Picou & Ricketts, 2014). In the Framework for Understanding Effortful Listening (FUEL), Pichora-Fuller and colleagues (2016) adapted a capacity model first proposed by Daniel Kahneman (1973) regarding attention under the lens of listening effort. They furthered the adapted model, offering a function of effort contingent on the task demands (in our context, background noise) and the motivation the person has to engage in the task. Although vaguely classified in the model, motivation is commonly defined as extrinsic or intrinsic, where both influence performance during a task. Extrinsic motivators, commonly referred to as incentives, are used in research tasks and real-world paradigms as they can be much more motivating for participants and employees e.g., money/bonuses or positive feedback. Intrinsic motivation comes from an inherent internal drive or enjoyment independent of outcomes.

Effortful behaviors often require more energy—either physically or mentally, which may be metabolically costly and generally fatiguing. To measure this exertion, Heitz et al., (2008) demonstrated motivation increased effort and performance during a working memory task.

Working memory typically requires variable levels of mental effort depending on the demands imposed and is often related to and considered as a factor during studies of effortful listening, making them suitable tasks to measure motivational effects (Rönnberg et al., 2013). Furthermore, Heitz and colleagues (2008) found that when separating a group of participants by their working memory capacity (high vs. low) and introducing within these group different levels of incentives (no incentives, feedback, feedback & money) they found that with more incentives, task performance was enhanced and evoked a larger pupil dilation response, a measure of effort.

However, even the highest incentivized, low-working memory group failed to perform better than the no-incentive, high-working memory group even though the former seemed to demonstrate more effort as assessed physiologically. This and other studies therefore suggest an important role of motivation on cognitive performance (Heitz et al., 2008, Ahern & Beatty, 1979; Hendijani et al., 2016; Herlambang et al., 2021; van der Meer et al., 2010; Zekveld et al., 2018).

According to various theories, extrinsic motivators may demonstrate an undermining effect leading to a decrease in intrinsic motivation where incentives may displace motivational salience towards obtaining a reward of the task rather than on the performance within a task (Deci, 1971; Deci et al., 1999). Contrarily, incentives and intrinsic motivation may show an additive effect where when used together, incentives enhance intrinsic motivation thus improving performance and overall motivation (Covington & Müeller, 2001). A study by Hendijani and colleagues (2016) set out to explore the interaction between incentives and intrinsic motivation by testing several hypotheses relating to the potential undermining and additive effects on performance. These researchers used a unique approach to intrinsic motivation by recruiting participants based on their chosen college major—math and literature, respectively. They then administered math and literature problems that would allow participants to earn incentives based on performance. With the hypothesis that mathematics majors would be more intrinsically motivated by mathematics problems and less by literature problems, and vice versa, the researchers found support for the additive effect of motivators on performance (Hendijani et al., 2016).

Certain social contexts are unique in that they may share both intrinsic and extrinsic motivational properties and the associated additive effect (Ryan & Deci, 2000). In these contexts, prosociality is a type of social motivation that is experienced as the drive to help and share with others and individuals with high social orientation have been shown to find value through cooperation, empathy, and other prosocial behaviors (Penner et al., 2005; Amani, 2022; Pfattheicher et al., 2022; Li et al., 2024). Social interactions are rewarding (Krach et al., 2020) and prosocial behaviors are often key components to these rewarding social interactions (Penner et al., 2005). An example of prosocial behavior was explored where Lockwood et al., (2021) tested whether age played a determining factor in how an individual would grip on a device in social interactions to either reward themselves or others. As opposed to the younger group, they found older individuals tended to exert more grip for others but also showed less self-bias for themselves (see also Li et al., 2024). As such, prosociality and other important social factors such as age and identity may be enhancing the integration of motivation that individuals use to physically and mentally engage in during social exchanges (Krach et al., 2010).

### C. Social Traits and SPiN

Noisy environments involve acoustic aspects that place demands on signal-level processing and cognitive and perceptual processing. As such, particular individual differences may underly variations in processing ability. For example, age is an influential factor often associated with general decreased hearing ability as well as decreased cognitive capacities leading to increased listening effort (Peelle et al., 2010; Degeest et al., 2015; Kwak & Han, 2021). However, there are few studies that address how social-psychological individual differences—such as prosocial orientation, affect SPiN experiences.

In one of the few studies looking at social traits and SPiN, Wöstmann et al., (2021) focused on personality and how it may be associated with performance and subjective experience during a hearing-in-noise task. Evidence from their study showed individuals who demonstrated higher neuroticism and lower extraversion tended to exhibit reduced self-ratings in respect to their listening performance. Neuroticism is the personality trait associated with negative emotions e.g., anxiety, frustration, jealousy, etc. and extraversion is associated with seeking and engaging in social interactions—opposed to introversion which is related to being more concerned with oneself than others. Most notably, higher neuroticism had positive associations with listening effort and performance during the SPiN task, albeit small effects. There was, however, only constant steady-state noise was used; therefore, the findings may only be generalizable to conditions related to pure energetic/modulation masking. Overall, Wöstmann et al. (2021) demonstrated individual differences regarding personality traits had effects on both performance and subjective listening effort during a SPiN task. Thus, there seems an underlying aspect of personality and the role of self-doubt, confidence, and modesty arising from these findings when thinking about a listener’s ability to perform in SPiN.

Additionally, Amani (2022) was interested in the moderating effects of emotional intelligence (EQ) on personality and found EQ played a key role in prosocial outcomes. Furthermore, EQ showed a moderating effect that proved to have the largest effect on the prosocial behavioral outcomes as well as having significant interactions with the three specific personality traits—neuroticism, agreeableness, and openness. Prosocial behavior is proposed to increase social support and self-efficacy during highly demanding listening situations, which are two stigmatized features common to SPiN, especially in older populations (Pichora-Fuller, 2016; Pfattheicher et al., 2022). Therefore, with a link between prosocial behavior and personality, and an association between personality and SPiN, there may be underlying aspects of prosociality that influence SPiN performance and listening effort that are yet to be explored and elucidated.

### D. Aims & Hypotheses

Motivation affects effort and fatigue during tasks, but the role of prosociality on listening effort and performance remains unclear. The main aim of this study was to take a novel approach to SPiN by relating performance and listening effort to scales regarding social reward and prosociality. The logic behind using prosociality to probe motivation is due to the idea that listening to someone speak innately requires cooperation and turn-taking via the sharing between a speaker and a listener. Although not the main focus of the study, we also included a measure of working memory capacity as working memory is an important underlying faculty in language processing (Rönnberg et al., 2013). Thus, we aimed to replicate findings pertaining to SPiN and working memory as well as explore any relationship between working memory and prosociality.

Individuals with increased social orientation tend to demonstrate more empathetic, kind, and reciprocal social behaviors (Lockwood et al., 2017) and may be more likely to perform listening tasks with more effort. Therefore, we hypothesized that individuals who self-rate higher on social reward and prosocial items will also demonstrate better performance in our SPiN task. Lastly, we hypothesized that listening effort will relate to different levels of motivation. Since task demands are increased in more adverse listening conditions, increased listening effort ratings may be associated to higher ratings of prosociality.

## III. Methods

### A. Participants

136 participants were recruited from the greater London area, UK using several recruitment channels including a university subject pool (SONA), word of mouth, and online advertisements. Exclusion criteria included the use of hearing aids, clinically significant hearing impairment, or a diagnosis of neurological or psychiatric disorders. Participants were self-reported native speakers of English or at bilingual-level proficiency. Several participants reported being proficient in or speaking two or more languages, including English (n = 65).

#### Stimuli

### B. Children’s Coordinate Response Measure (CCRM)

The CCRM is a corpus of sentences used to assess speech perception—similar to the coordinate response measure (Bolia et al., 2000) and first introduced by Messaoud-Galusi et al. (2011).

Three speech masking conditions were used, 1) a steady-state speech-spectrum-shaped noise (SSN), 2) a 1-speaker condition, and 3) a 2-speaker condition. The SSN condition acts as a purely energetic masker whereas the two maskers with competing speakers are primarily informational maskers. The target speaker and competing speakers are different native British adult males. The target speaker was randomly one of three male potential speakers. Additionally, the competing speakers were never the same as the target speaker, nor the other masking speaker. The target speech in all conditions and trials directs the participant to the target speaker with the call sign “*Show the dog where the [color] [number] is*”, requiring the participant to select the corresponding color-number pair. Background speech was sentences unrelated to the target speech. The color-number combinations are random and included any combination of six monosyllabic colors (red, white, blue, green, pink, black) and eight numbers (1-9, excluding 7). If a participant responded accurately by selecting the correct combination, a happy bear is presented as feedback; if incorrect, a sad bear.

This task adaptively determines at what SNR listeners will accurately respond to the target speech approximately 50% of the time, the so-called speech reception threshold (SRT). This speech-in-noise task was administered with MATLAB R2022b, and all participants listened to the sounds at the same comfortable sound pressure level (SPL) through Sennheiser HD221 headphones, with the option to adjust the volume manually on the computer if that felt necessary. Target and masker audio were simultaneously presented diotically through the headphones in non-sound-attenuating testing rooms.

The initial signal-to-noise ratio (SNR) is set at +12 dB, resulting in easily intelligible speech. The task follows a stepwise adaptive process based on the listener’s responses. For the first steps—defined by correct and incorrect responses—the SNR is adjusted by -9 dB, followed by +7 dB, -5 dB, and finally ±3 dB. Once the ±3 dB step is reached and speech becomes significantly less intelligible, a mean score from the last six reversals is calculated to determine the listener’s SRT. The test terminates after the sixth reversal or after 30 trials

### C. Reading Span Task

This cognitive task utilized an open-access, computerized version of the reading span task (RST; Daneman & Carpenter, 1980) where rather than repeating back words at the end of sentences in varying set sizes as the classic version did, participants recalled numbers after determining whether sentences make sense or not. This version was acquired from http://www.cognitivetools.uk/cognition/ (Stone & Towse, 2015) and administered on a Dell laptop using Tatool via JAVA (von Bastian et al., 2013). The RST measures working memory capacity (WMC) or the ability to process new information whilst holding previous bits of information in mind for recall. The information stored and recalled in each trial was a random number between 1-99. After the random number was presented in the center of the screen for 250 ms, a short sentence is presented and the participant processed and determined whether this sentence made sense or not using the keyboard’s arrow keys. Feedback is provided in the form of a green check for correct responses or a red ‘x’ for incorrect responses. The judgement is strictly based on semantics e.g. “*you can hear with your ears*” (sensical) or “*boats like to eat crisps*” (non-sensical). Following the sentences was a screen that prompts the participant to input the numbers previously presented before the sentences. The same feedback system is used for correct or incorrect number inputs. After each trial there is a 2500 ms pause before the next trial begins. The span length of numbers to remember ranges from two to six. There were three trials per span condition within the five different span lengths, equating to 15 total trials and a total of 60 number and 60 sentence responses. A working memory score was derived by taking the total number of correct number responses out of 60. Sentence accuracy was also calculated by total number of correct responses out of 60.

### D. Social Reward Questionnaire (SRQ)

The SRQ is a 20-item questionnaire that measures how rewarding or enjoyable individuals find various social interactions (Foulkes et al., 2014). The scale is divided into 5 subscales of social reward—Admiration, Negative Social Potency, Passivity, Prosocial Interactions, and Sociability. Items are scored using a 7-point Likert scale with points ranging from Strongly Disagree to Strongly Agree. Participants were instructed to report how much they imagine they would enjoy an item in the case they had never experienced that prompt.

According to Foulkes et al., (2014) each subscale—or factor, can be described as follows:

- Admiration: “*being flattered, liked and gaining positive attention*”
- Negative social potency: “*being cruel, callous and using others for personal gains*”
- Passivity: “*giving others control and allowing them to make decisions*”
- Prosocial interactions: “*having kind, reciprocal relationships*”
- Sociability: “*engaging in group interactions*”

The SRQ subscales are unidimensional and were subsequently collapsed into five individual aggregate scores.

### E. Prosocial Scale for Adults (PSA)

The PSA is a 16-item questionnaire that measures differences in prosocial tendencies (Caprara et al., 2005). The scale instructs participants to mark their first reaction to different, common intersocial scenarios. Self-rated items e.g., “*I easily help others*” get scored on a 5-point Likert scale where values represent Never/Almost Never, Rarely, Occasionally, Often, and Always/Almost Always. This scale is considered unidimensional; therefore, all item scores were collapsed into one aggregate score for prosociality.

## IV. Procedure

Participants were tested onsite at University College London, Institute of Cognitive Neuroscience (UCL ICN). Informed consent and demographic information were obtained from each participant following the approval from the UCL Research Ethics Committee. Participants first completed the Children’s Coordinate Response Measure (CCRM)—after each condition, participant were asked to rate their subjective listening experiences on the CCRM questionnaire.

These ratings pertained to the noise they experienced, the effort they exerted, and how aware of their effort they were, all between 0 and 10. The three masker conditions were pseudorandomized to eliminate any potential learning effects in performance and adaptations in subjective reports. Participants then completed the reading span task (RST) and lastly, the paper-and-pen SRQ and PSA. After all tasks and scales were administered, participants were allowed to ask any questions and provided a debrief regarding the aims of the study. All participants were compensated 10 British Pound Sterling for their time and efforts.

Prior to the analyses, a principal component analysis (PCA) was performed on the scores produced by the social reward and prosocial questionnaires (FactoMineR; Lê et al., 2008). PCA was used to reduce the dimensionality of these metrics in order to elucidate any underlying structure amongst these variables. PCA indicated there were two principal components that accounted for 60.6% of the total variance (40.3% and 20.3%, respectively). The first component (PC1/prosocial reward) was most strongly influenced by prosociality (PSA), prosocial interactions, sociability, negative social potency and admiration; suggesting this dimension of social reward represents prosocial social traits. The second component (PC2/passive reward) was most strongly influenced by passivity, negative social potency, admiration, and sociability; suggesting a component of social reward represented by passive social traits. These two dimensions represent a key difference pertaining to the moderating role of prosocial motivation and passivity in social reward. Therefore, these two components are used as variables to explore relationships between prosociality and objective and subjective SPiN. Moreover, due to a high correlation between subjective ratings in all conditions post-CCRM, these responses were aggregated as one single metric for each condition which will be referred to as *listening effort*.

Data was primarily analyzed with JASP and RStudio. All reported statistics have undergone assumption checks and if any assumptions were violated, corrected statistics are reported when applicable. Correlation p-values were corrected for multiple comparisons using the false discovery rate (FDR) correction (Benjamini & Hochberg, 1995). Linear mixed models (LMM) were utilized to address the issue of unequal sample sizes, particularly between masker conditions in the CCRM. There were also few instances of highly influential outlying data points. Data outside of three standard deviations from the mean were excluded from analyses. Analyses were performed on the outcomes from SRTs and listening effort. The models had fixed effects of masker condition, age, sex, reading span scores, and PCA components and a random effect of participant ID.

For regression, outliers were labelled as negatively influential according to a relatively conservative Cook’s distance (Cook’s d = 0.1). Any outliers with a Cook’s d larger than 0.1 were subsequently removed from the analyses. Furthermore, backwards elimination stepwise regression was used to elucidate meaningful relationships between outcome and predictor variables. Backwards elimination stepwise regression considers all predictor variables and subsequently eliminates each variable from the model one-by-one starting with the least insignificant (highest p-value) variable until there only remains a model—or models, with significant predictors (threshold of p = 0.05).

## V. Results

Of the 136 participants, there were 83 female and 53 male. Mean age was 29.6 years (SD = 10.6, range = 18-68). Male participants were significantly older (M = 33.2, SD = 9.1) than female participants (M = 27.4, SD = 12; t(133) = -3.189, p = .002). One male participant failed to provide age information.

## A. SPiN Performance & Listening Effort

Linear mixed-effects models (LMM) were used to explore the differences between the masker conditions in the CCRM in SRTs and the subjective ratings of effort. There was a significant main effect of masker on SRT (F[2, 261.54] = 734.66, p < .001; Figure 1) and a main effect of prosocial reward (F[1,125.67] = 8.549, p = .004; Table 1). As for subjective listening effort, there were significant main effects of age (F[2,127.02] = 5.5, p = .021) and masker (F[2,263.31] = 15.25, p < .001; Table 2). SRTs were poorest in the 2-speaker masker condition, which also elicited the highest self-ratings of listening effort (Figure 2). Post-hoc comparisons (Table 3 & 4) show that the SRTs were significantly different from each other in all maskers whereas subjective listening effort ratings were only significantly different between the 1-speaker and 2-speaker maskers.

**Figure 1:**
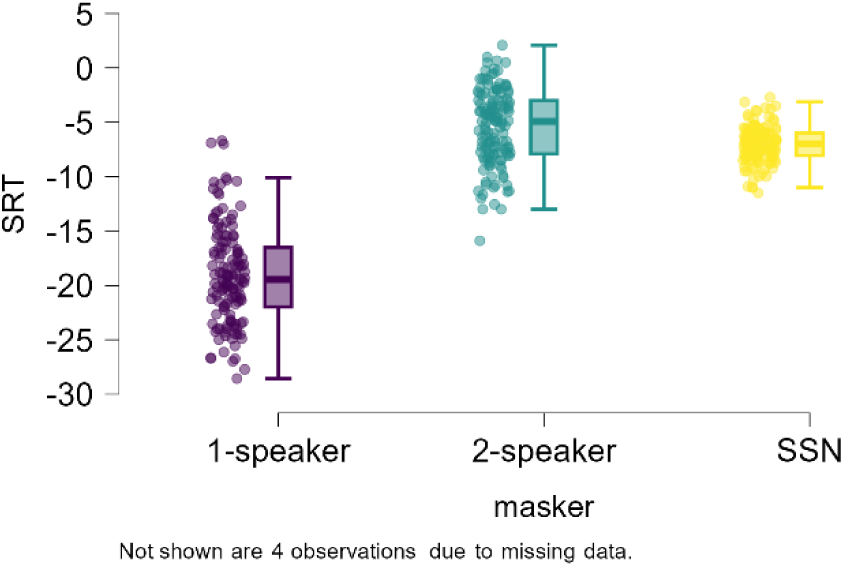
right) boxplots showing mean SRTs for each masker condition.

**Figure 2:**
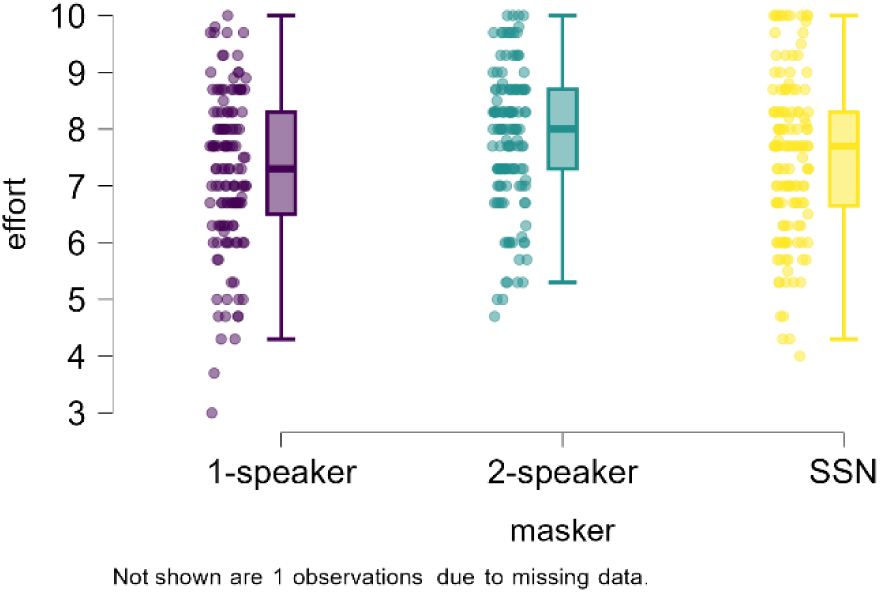
right) boxplots for the subjective ratings of listening effort for each condition.

**Table 1:**
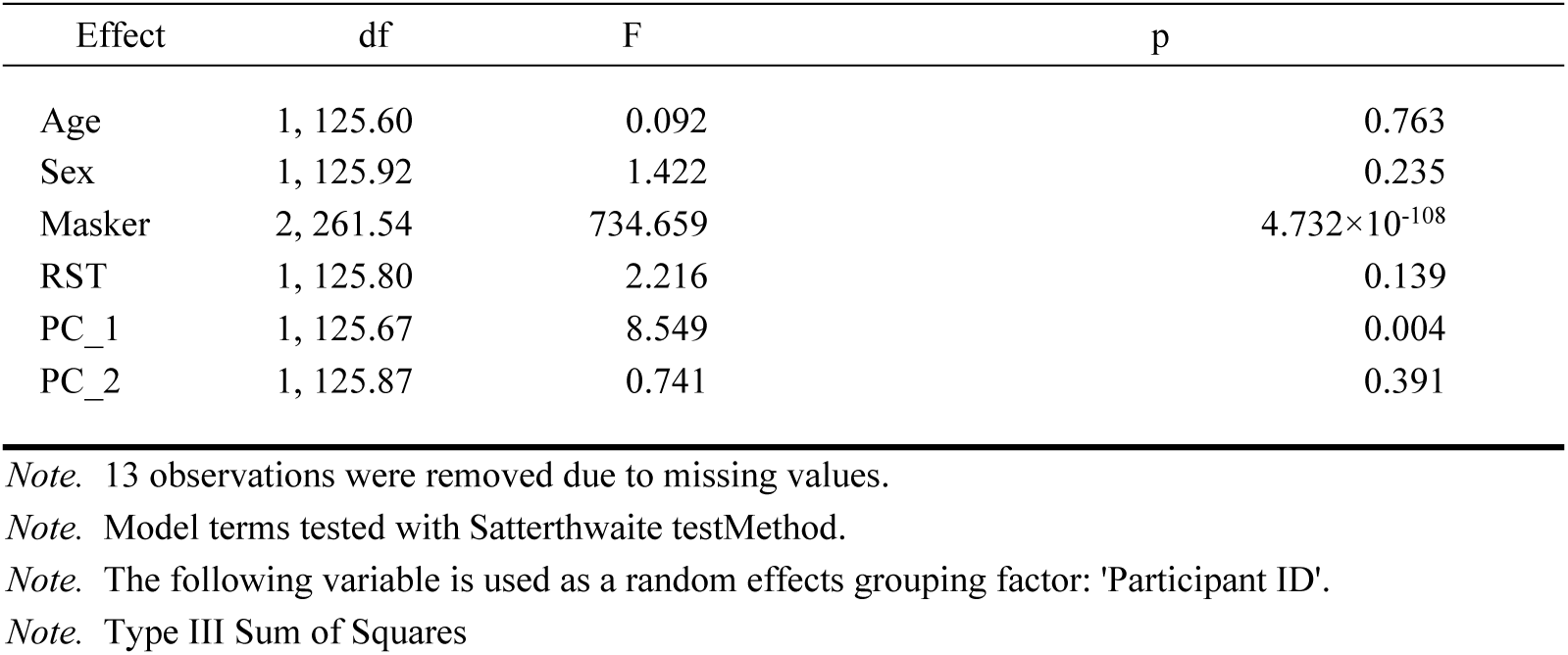
ANOVA Summary.

| Effect | df | F | p |
| --- | --- | --- | --- |
| Age | 1, 125.60 | 0.092 | 0.763 |
| Sex | 1, 125.92 | 1.422 | 0.235 |
| Masker | 2, 261.54 | 734.659 | $4.732 \times 10^{-108}$ |
| RST | 1, 125.80 | 2.216 | 0.139 |
| PC_1 | 1, 125.67 | 8.549 | 0.004 |
| PC_2 | 1, 125.87 | 0.741 | 0.391 |

Note. 13 observations were removed due to missing values.
Note. Model terms tested with Satterthwaite testMethod.
Note. The following variable is used as a random effects grouping factor: 'Participant ID'.
Note. Type III Sum of Squares

**Table 2:** ANOVA Summary.

| Effect | df | F | p |
| --- | --- | --- | --- |
| Age | 1, 127.04 | 5.956 | 0.016 |
| Sex | 1, 126.96 | 1.862 | 0.175 |
| Masker | 2, 263.29 | 15.253 | $5.396 \times 10^{-7}$ |
| RST | 1, 127.01 | 0.031 | 0.861 |
| PC_1 | 1, 127.14 | 0.251 | 0.617 |
| PC_2 | 1, 127.08 | 0.127 | 0.722 |

Note. 10 observations were removed due to missing values. Note. Model terms tested with Satterthwaite testMethod. Note. The following variable is used as a random effects grouping factor: 'Participant ID'. Note. Type III Sum of Squares

**Table 3:**
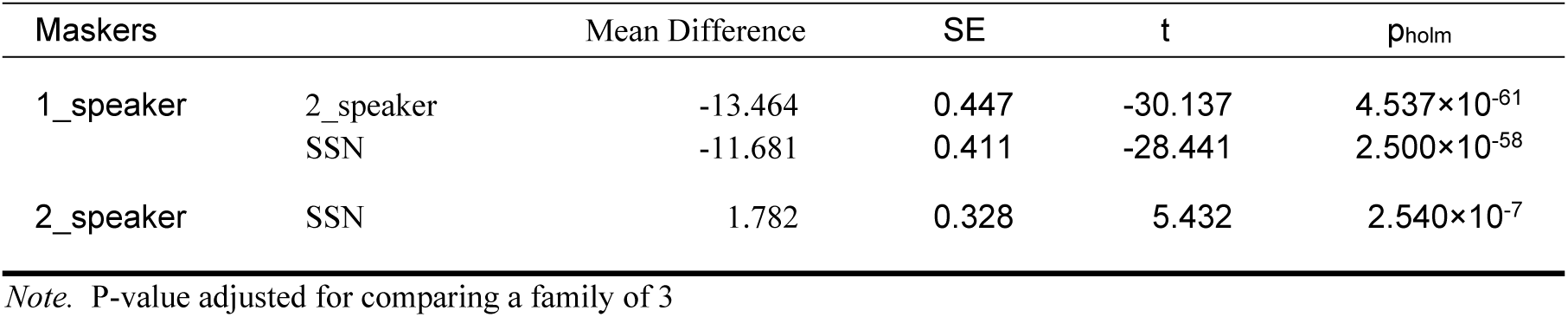
Post Hoc Comparisons - SRTs.

### B. Relationships in SPiN

Correlation and regression models were used to explore relationships between predictor variables (age, sex, working memory, and PCA components) and outcome variables (performance and listening effort). Figure 3 shows a correlation heatmap of variables. Of the correlations that survived FDR correction, the most notable were between SRTs in the 2-speaker condition and the prosocial reward component (p_FDR_ = .018), between age and the passive reward component (p_FDR_ = .005), and between age and listening effort in the 1-speaker condition (p_FDR_ = .039). All listening effort ratings showed intercorrelation between masker types, whereas the only relationship between SRTs that survived FDR correction was between the 1-speaker and SSN maskers. No variables related to working memory scores, therefore we failed to replicate previous findings pertaining to the relationship between working memory and SPiN.

**Figure 3:**
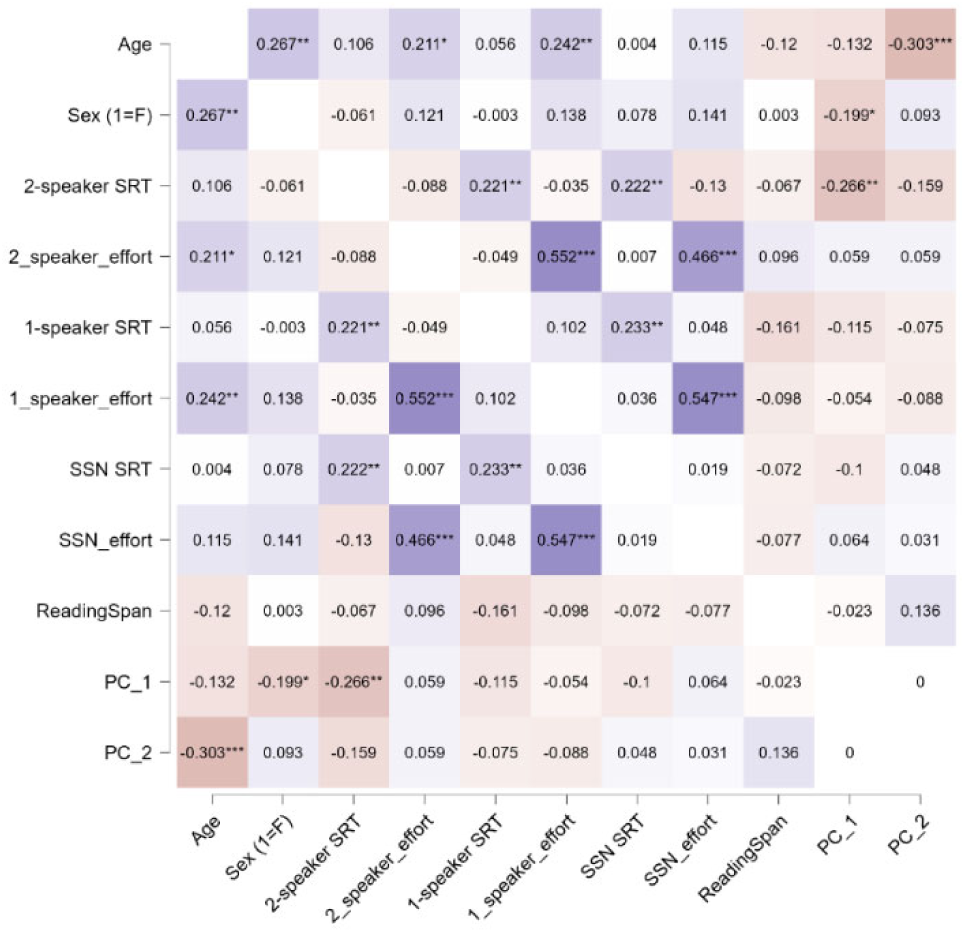
Pearson r heatmap of all variables. * p < .05, ** p < .01, *** p < .001 (uncorrected)

As for regression, the model that demonstrated the best predictive power for SRTs (F(2,392) = 195.91, p < .001; Table 5), accounting for 49.7% of the variance, was influenced by the masker condition and the component of prosocial reward (Figure 4).

**Figure 4:**
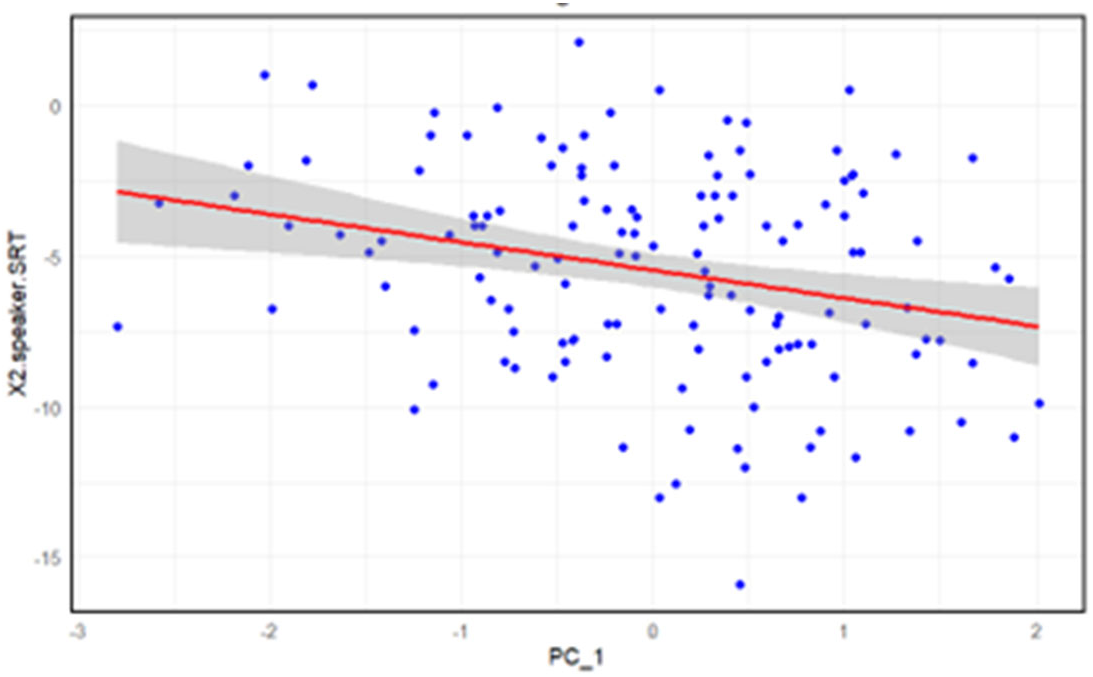
Negative relationship between prosocial reward and SRTs, where higher prosocial reward is associated with better SRTs in the 2-speaker masker.

**Table 4:** Post Hoc Comparisons – Listening Effort.

| Maskers |  | Mean Difference | SE | t | p<sub>holm</sub> |
| --- | --- | --- | --- | --- | --- |
| 1_speaker | 2_speaker | -0.587 | 0.105 | -5.577 | 3.892×10<sup>-7</sup> |
|  | SSN | -0.211 | 0.113 | -1.863 | 0.065 |
| 2_speaker | SSN | 0.376 | 0.116 | 3.224 | 0.003 |

Note. P-value adjusted for comparing a family of 3

**Table 5:** SRT Coefficients.

| Model |  | Unstandardized | Standard Error | Standardized | t | p |
| --- | --- | --- | --- | --- | --- | --- |
| M<sub>0</sub> | (Intercept) | -21.047 | 1.408 | | -14.944 | $3.329 \times 10^{-40}$ |
|  | Age | 0.006 | 0.026 | 0.009 | 0.222 | 0.825 |
|  | RST | -1.747 | 1.483 | -0.043 | -1.178 | 0.240 |
|  | Sex (1=F) | -0.540 | 0.551 | -0.038 | -0.980 | 0.328 |
|  | PC_1 | -0.604 | 0.261 | -0.084 | -2.311 | 0.021 |
|  | PC_2 | -0.182 | 0.271 | -0.026 | -0.670 | 0.503 |
| | masker | 6.017 | 0.306 | 0.702 | 19.646 | $3.797 \times 10^{-60}$ |
| M<sub>1</sub> | (Intercept) | -20.910 | 1.262 | | -16.564 | $5.144 \times 10^{-47}$ |
|  | RST | -1.777 | 1.476 | -0.043 | -1.204 | 0.229 |
|  | Sex (1=F) | -0.505 | 0.527 | -0.035 | -0.958 | 0.339 |
|  | PC_1 | -0.608 | 0.260 | -0.085 | -2.333 | 0.020 |
|  | PC_2 | -0.201 | 0.257 | -0.029 | -0.784 | 0.433 |
| | masker | 6.017 | 0.306 | 0.702 | 19.670 | $2.721 \times 10^{-60}$ |
| M<sub>2</sub> | (Intercept) | -20.762 | 1.248 | | -16.641 | $2.284 \times 10^{-47}$ |
|  | RST | -1.929 | 1.462 | -0.047 | -1.319 | 0.188 |
|  | Sex (1=F) | -0.558 | 0.522 | -0.039 | -1.069 | 0.286 |
|  | PC_1 | -0.604 | 0.260 | -0.084 | -2.321 | 0.021 |
| | masker | 6.018 | 0.306 | 0.702 | 19.682 | $2.195 \times 10^{-60}$ |
| M<sub>3</sub> | (Intercept) | -21.522 | 1.025 | | -20.992 | $4.608 \times 10^{-66}$ |
|  | RST | -1.941 | 1.462 | -0.047 | -1.327 | 0.185 |
|  | PC_1 | -0.554 | 0.256 | -0.077 | -2.164 | 0.031 |
| | masker | 6.016 | 0.306 | 0.702 | 19.673 | $2.181 \times 10^{-60}$ |
| M<sub>4</sub> | (Intercept) | -22.563 | 0.661 | | -34.132 | $1.839 \times 10^{-119}$ |
|  | PC_1 | -0.547 | 0.256 | -0.076 | -2.133 | 0.034 |
| | masker | 6.019 | 0.306 | 0.702 | 19.665 | $2.148 \times 10^{-60}$ |

The regression model providing the best fit for listening effort (F(2,395) = 10.49, p < .001; Table 6), accounting for 4.6% of the variance in the model, was primarily influenced by age (Figure 5) and by the sex of the participant.

**Figure 5.**
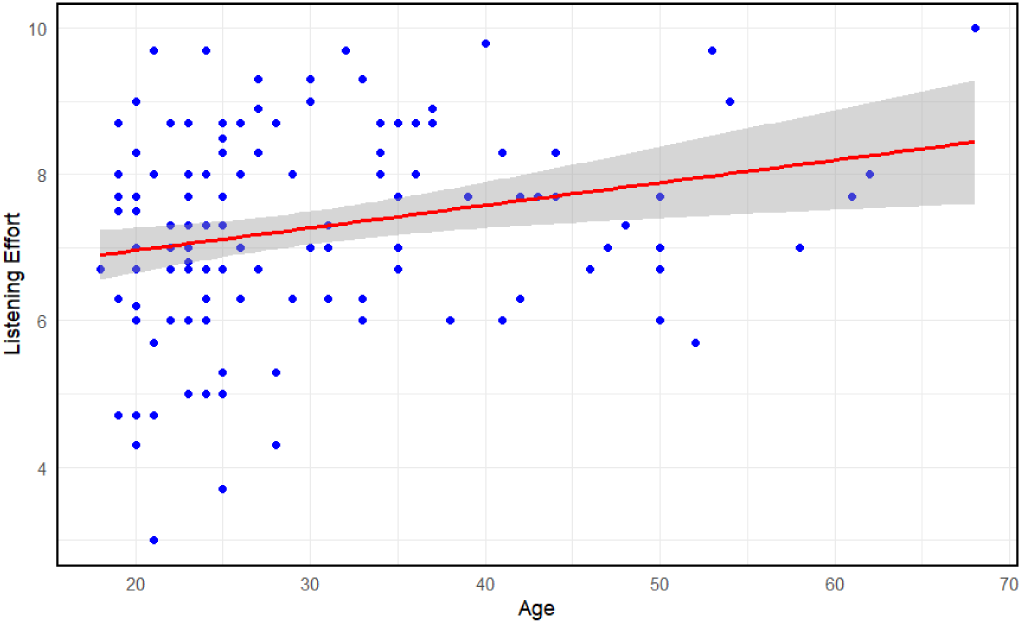
positive relationship between age and the subjective effort in the 1-speaker masker.

**Table 6:** Listening Effort Coefficients.

| Model |  | Unstandardized | Standard Error | Standardized | t | p |
| --- | --- | --- | --- | --- | --- | --- |
| M<sub>0</sub> | (Intercept) | 6.248 | 0.375 | | 16.679 | $1.482 \times 10^{-47}$ |
| | Age | 0.024 | 0.007 | 0.187 | 3.453 | $6.141 \times 10^{-4}$ |
|  | Sex (1=F) | 0.283 | 0.146 | 0.102 | 1.933 | 0.054 |
|  | RST | -0.093 | 0.394 | -0.012 | -0.237 | 0.813 |
|  | PC_1 | 0.050 | 0.070 | 0.036 | 0.723 | 0.470 |
|  | PC_2 | 0.037 | 0.072 | 0.027 | 0.517 | 0.606 |
|  | masker | 0.128 | 0.081 | 0.077 | 1.578 | 0.115 |
| M<sub>1</sub> | (Intercept) | 6.194 | 0.298 | | 20.762 | $4.005 \times 10^{-65}$ |
| | Age | 0.024 | 0.007 | 0.188 | 3.494 | $5.310 \times 10^{-4}$ |
|  | Sex (1=F) | 0.282 | 0.146 | 0.101 | 1.932 | 0.054 |
|  | PC_1 | 0.051 | 0.069 | 0.036 | 0.729 | 0.466 |
|  | PC_2 | 0.036 | 0.072 | 0.026 | 0.497 | 0.619 |
|  | masker | 0.128 | 0.081 | 0.077 | 1.580 | 0.115 |
| M<sub>2</sub> | (Intercept) | 6.208 | 0.297 | | 20.919 | $7.597 \times 10^{-66}$ |
| | Age | 0.023 | 0.006 | 0.179 | 3.534 | $4.580 \times 10^{-4}$ |
|  | Sex (1=F) | 0.298 | 0.143 | 0.107 | 2.088 | 0.037 |
|  | PC_1 | 0.049 | 0.069 | 0.035 | 0.711 | 0.478 |
|  | masker | 0.128 | 0.081 | 0.077 | 1.580 | 0.115 |
| M<sub>3</sub> | (Intercept) | 6.240 | 0.293 | | 21.281 | $1.867 \times 10^{-67}$ |
| | Age | 0.023 | 0.006 | 0.177 | 3.504 | $5.118 \times 10^{-4}$ |
|  | Sex (1=F) | 0.281 | 0.141 | 0.101 | 2.000 | 0.046 |
|  | masker | 0.128 | 0.081 | 0.077 | 1.579 | 0.115 |
| M<sub>4</sub> | (Intercept) | 6.496 | 0.245 | | 26.564 | $6.319 \times 10^{-90}$ |
| | Age | 0.023 | 0.006 | 0.177 | 3.495 | $5.280 \times 10^{-4}$ |
|  | Sex (1=F) | 0.281 | 0.141 | 0.101 | 1.993 | 0.047 |

## VI. Discussion

We first tested the hypothesis that individuals who rate themselves higher in prosociality perform better SRTs during a SPiN task. In this case, we rejected the null hypothesis as novel findings from the study show under certain masker conditions e.g., high-difficulty speech-on-speech conditions, people who rated themselves higher in prosocial reward demonstrated better SRTs, particularly the 2-speaker condition. We also test the hypothesis whether individuals would vary in listening effort based on these motivational traits. We failed to reject the null hypothesis, as age proved to be the primary predictor of self-rated listening effort during the SPiN task. Lastly, we also failed to replicate previous findings related to the relationship between working memory and SPiN performance.

It is important to note that we did not collect any audiometric data, and the SPiN task is the only measure of hearing ability in this study however, there were no cases that would have led us to believe a participant exhibited any hearing difficulties. We also included age and sex as potential factors influencing performance and found no main effects or interactions between these variables. Given the limited hearing ability information, the masker-effect independent of age and sex aligns with previous research in that informational maskers and energetic maskers impose variations of demands on different levels of processing (Shinn-Cunningham, 2008; Scott et al., 2004; Evans et al., 2016, Brungart, 2001).

### A. Speech Masking Noise Effects

More specifically, background noises that are amplitude modulated provide increased chances for listeners to “glimpse” target speech (Brungart, 2001; Mattys et al., 2009; Scott et al., 2004). Glimpsing target speech is directly associated with specific acoustical features, particularly the number of background speakers (Cooke et al., 2008; Zhang et al., 2021). Rosen and colleagues (2013) demonstrated that a condition with 2-speaker babble proved the most difficult in terms in accuracy performance; and that, when incrementally increasing the number of competing speakers up to 16 or infinity, there was only a small increase in performance. This is because the amount of information that can be accessed from each speaker diminishes as aperiodic speech streams overlap one another, leading to less glimpsing and increased energetic masking (Steinmetzger & Rosen, 2015).

In our case, the masker condition in this study with two speakers competing against target speech resulted in the worst thresholds out of the three conditions. The condition with only one speaker yielded the best SRTs. However, the SSN condition (infinity speakers) consistently elicited SRTs that were closer to the 2-speaker condition yet demonstrated a slight, yet significant improvement in thresholds. This finding highlights an underlying nonmonotonic relationship between the number of speakers and masking effects, and this relationship is likely the reason behind the robust difference between SRTs found in our study.

### B. Social Traits and SPiN

We also found evidence through a significant relationship between performance and prosocial reward that individuals who self-rated themselves as more prosocial tended to perform with better SRTs in the 2-speaker condition. This finding fills an important gap by linking social motivational traits to SPiN performance, particularly within the framework proposed by Pichora-Fuller et al. (FUEL; 2016). However, one major limitation of the current study is the independence between measures; therefore, the correlational findings may not allow for strong generalizable conclusions, leaving us to simply infer how prosociality plays a motivational role during our SPiN task.

Given this inference, one explanation for this finding is that more prosocial individuals may navigate noisy conditions with increased confidence and better subsequent performance in these conditions (Pichora-Fuller, 2016). It may be that increased prosociality increases affinity for social environments, leading to more exposure and experience for dealing with adverse listening environments. Through increased experience and having strong beliefs in one’s listening abilities, enjoying social settings and providing support to others during noisy situations becomes a rewarding interaction where both the individual and the other benefit through combinations intrinsic and extrinsic properties. Therefore, prosocial behavior may aid in adverse conditions by encouraging and rewarding interactions amidst noise, which then helps listeners gain experience and techniques for efficient mental resource allocation during SPiN.

Our study included the SRQ which contained a subscale of sociability—a strong factor in our prosocial reward component, indicating pleasure in group settings is related to prosociality and cocktail party listening performance. In most SPiN literature, there is a primary focus on general hearing ability, cognitive abilities, and perceptual skills related to phoneme and word discrimination. However, Foulkes et al. (2014) found prosociality to be highly correlated with personality whereas Wöstmann et al. (2021) noted relationships between personality and hearing-in-noise. So based on the findings of the current study, it seems feasible to consider and include psycho-social factors such prosociality and personality when investigating individual differences during SPiN in the future.

### C. Age and Effort

There was an age difference between the male and female groups, however no notable sex differences were found in any of the measures. Although there was no relationship to age and performance, the noted age effect on effort in the 1-speaker condition is consistent not only with previous SPiN studies (Koelewijn et al., 2012) but previous literature investigating listening effort in older adults. Older adults tend to face higher demands on resource allocation to accurately perform, often resulting in more subjective effort due to decreased mental capacities such as memory, attention, etc. (Peelle et al., 2010; Degeest et al., 2015; Kwak & Han, 2021).

Moreover, individuals often express more social effort as result of increased prosociality across their lifespan (Lockwood et al., 2021; Li et al., 2024). Therefore, it is important to consider that although individuals may increasingly orientate socially towards others as they get older, they may also experience reduced social connectedness and impaired situational awareness in parallel (Kwak & Han, 2021), which are key social communication factors linked to belonging, stress, and performance (Pichora-Fuller, 2016). Additionally, higher social connectedness was found to be associated with increased listening effort in a sample of cochlear implant users (Hughes et al., 2018). Similarly, other studies looked at the size of social networks and the relation to speech perception abilities, but yielded mixed results (Lev-Ari, 2018; Lev-Ari, 2022).

Furthermore, age was negatively associated to our second PCA component—passive reward. As mentioned previously, passivity is referred to as giving others control and allowing them to make decisions, and although this was not related to listening effort, it seems that older people are motivated to have more control and decision-making abilities in social environments compared to younger people, a seemingly opposite construct to that of prosociality.

## VII. Limitations

As mentioned above we failed to conduct audiometry, meaning there may have been instances of untested age-related or other forms of hearing loss given the large range of age in our participants; however, there were no major outlining data points in the CCRM that would suggest any of our participants had significant hearing difficulties. Moreover, our speech in noise task was independent of the social trait questionnaires, leaving most of the interpretations to inference and correlation. Future studies would benefit in listening paradigms that elicit some for of reciprocity, sharing, and cooperation beyond strict recognition in noise. We also failed to include any formal measure or inquiry of the participant’s fatigue. Inter-trial fatigue could have had effects on the effort ratings. Lastly, sample size can be increased. While we had a substantial sample size for our study, more participants may allow for greater detailed correlations of subtle individual differences.

## VIII. Conclusion

This study explored whether prosociality would influence listener’s performance and effort during a SPiN task, and is to our understanding, the first study to do so. The findings of this study provide novel insights into the relationship between prosociality and speech perception in noise performance as well as information pertaining to social orientations in older adults. We were also able to replicate findings pertaining to age and listening effort, but failed to find any relationship between working memory, performance, and listening effort. In total, these findings highlight the importance of considering psycho-social traits in understanding both objective performance and subjective listening experiences during speech perception. Further research should aim to explore the physiological and neurobiological mechanisms underlying prosociality and its impact on cognitive processing during challenging language-related tasks.

## Author Declaration

We, the authors, confirm that there are no known conflicts of interest associated with this publication and there has been no significant financial support for this work that could have influenced its outcome. The manuscript has been read and approved by all associated authors named.

## Data Availability

The data that support the findings of this study are openly available in OSF at https://osf.io/9gbfr/ or DOI 10.17605/OSF.IO/9GBFR.

## Ethics Approval

Ethical approval for this study was obtained from UCL Research Ethics Committee (approval ID number: ICN-VW-10-5-23A).

